# Multi-trait eco-evolutionary dynamics explain niche diversity and evolved neutrality in forests

**DOI:** 10.1101/014605

**Authors:** Daniel Falster, Åke Brännström, Mark Westoby, Ulf Dieckmann

**Affiliations:** Department of Biological Sciences, Macquarie University NSW 2109, Australia; Evolution and Ecology Program, International Institute for Applied Systems Analysis, Schlossplatz 1, A-2361 Laxenburg, Austria; Department of Mathematics and Mathematical Statistics, UmeÅ University, 901 87 Umeå, Sweden; International Institute for Applied Systems Analysis, Austria

## Abstract

An enduring challenge in ecology is to understand how diverse plant species coexist when competing for the same basic resources^1–4^. Two candidate frameworks for meeting this challenge currently exist in forest ecology. Niche-based approaches succeed in describing successional dynamics in response to recurrent disturbances^5–9^. Approaches based on Hubbell’s neutral theory describe abundance patterns in highly diverse communities^10^. Yet both theories are unsuccessful where their competitor is strong, highlighting an enduring need for reconciliation^11–13^. In fact, the perception that niche and neutral processes are incompatible may have arisen because both types of models have lacked some critical features of real forests. Here we report a productive reconciliation that arises from extending niche models to include three ubiquitous features of vegetation: (1) plant growth under light competition, (2) multiple trait-mediated tradeoffs in plant function, and (3) evolutionary community assembly. We show that fitness equivalence – which neutral theory controversially takes as an assumption^4,12,13–14^ – naturally arises as a mechanistic outcome of niche differentiation, but only within a specific region of trait space, corresponding to shade-tolerant later-successional strategies. In this way, niche diversification can lead, via evolved neutrality, to a proliferation of shade-tolerant strategies, one of the phenomena that motivated the development of neutral theory^12^. Earlier niche models focussed on a single growth-mortality trait and found no evolved neutrality. In contrast, the model introduced here includes a second trait, height at maturation, in addition to a growth-mortality trait in the form of leaf-construction cost. We show that fitness equivalence only emerges when diversifications include this second trait. We also demonstrate that our extended niche model generates markedly different forest types under different environmental conditions, recovering known biogeographic patterns^15–18^. By evolving forests from first principles, our study constructively resolves the protracted debate about niche versus neutral dynamics for forest communities, and provides a platform for a synthetic theory of forest structure and diversity.

## 2 MAIN TEXT

The central challenge of community ecology is to discern the rules governing the assembly of species and growth types. The problem is acute for vegetation, because differential resource utilization cannot explain the observed trait differences among plant species^1–3^. Yet most forests contain tens to hundreds of plant species^1,19–20,21^. Consequently, niche-based theories of biodiversity seek to identify mechanisms through which multiple species can coexist while competing for a single resource such as light^2,6,13,22–24^. Tradeoffs in plant function are key, because they limit one species from monopolising all available growth opportunities. In particular, a tradeoff between growth in high light and survival in low light has been shown theoretically to allow different successional types to coexist in vegetation subject to recurrent disturbance^23^. Data from tropical forests show that diverse assemblages are distributed along a single growth-mortality axis^10,25^ and thus support the potential importance of this tradeoff for community assembly. Species are distributed unevenly along this axis, with a proliferation of species clustered at the slow-growth-low-mortality end^10,19,25^. By contrast, existing niche models suggest that only a few species can be maintained at equilibrium, including only a single slow-growing species^5–9^. This mismatch between theory and observation has led many to question the importance of successional niche differentiation for community assembly and to seek alternative explanations of forest diversity^1,7,9,10,12,19^.

The unified neutral theory of biodiversity (UNT)^10^ offers such an alternative. Forest diversity is explained by ecological and evolutionary drift, via stochastic speciation and population dynamics^10,12^. Unlike niche theory, the UNT assumes that species are functionally identical. This implies that all individuals have equal per-capita birth and death rates, and thus are equivalent in their fitness. To most forest ecologists (including Hubbell himself^12^), this functional-equivalence seems implausible, partly for theoretical reasons^14^, but mainly because coexisting species are known to vary greatly in their traits and demographic rates^4,10,12,19^. Yet the UNT successfully captures features of tropical forests not explained by current niche theory, notably the substantial number of shade-tolerant species. It has been suggested that the UNT will continue to be supported as long as niche theory cannot *“come up with a plausible argument for why niches are more finely partitioned under low-light conditions than under high-light conditions”*^12^.

Here we show how a more realistic modelling of successional niche differentiation naturally leads to communities with a higher diversity of slow-growing strategies adapted to low-light conditions (Fig. 1). We extend classic niche models to include three ubiquitous features of vegetation. Although it is well known that each feature is important, none of them have previously been incorporated in niche models. First, we explicitly model the size-structured nature of growth, mortality, and light competition based on a well-established mechanistic understanding of growth and competitive shading (Fig. 1; Methods). This allows the fitness of plants to be evaluated over their entire life cycle^26^. Second, we move beyond general demographic tradeoffs by incorporating specific well-known tradeoffs in plant function. Species are represented by trait values indicating their position along these physiological tradeoffs. Demographic and growth differences are not assumed a priori, but naturally emerge from these trait differences (Fig. 6–7). Third, communities are assembled evolutionarily as well as ecologically. This allows us to assess how a current species mixture gives rise to a fitness landscape in trait space (Fig. 5). In turn, this fitness landscape determines the trait combinations that can invade, and hence the direction of evolutionary change. This approach enables us to capture the intricate dynamical feedback between a community’s fitness landscape and its composition. On this basis, we examine the trait mixtures expected to emerge through repeated rounds of immigration, mutation, and selection on an evolving fitness landscape. Together, these three advances allow us to investigate how multiple tradeoffs interact to shape forest diversity and structure.

**Figure 1:**
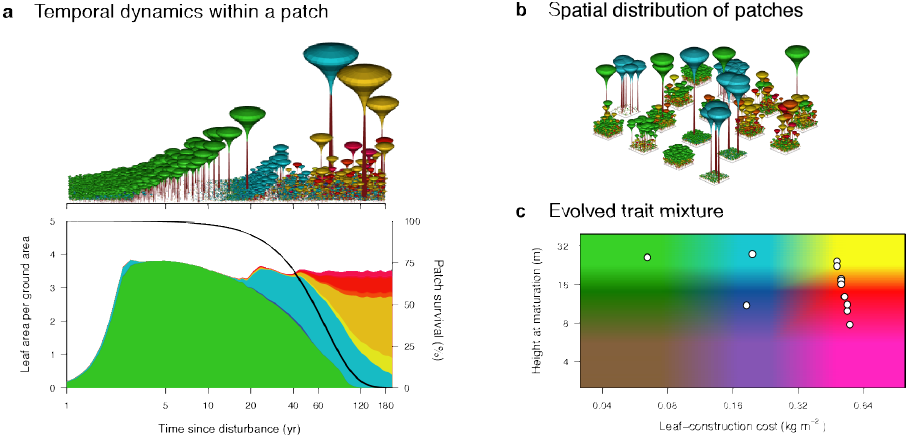
Multi-trait successional dynamics enable rich species coexistence. We consider a model in which plant species inhabiting a metapopulation of patches can differ in two functional traits, leaf-construction cost and height at maturation. a, After a disturbance, occurring on average every 60 years, vegetation development in a patch follows successional dynamics under size-structured competition for light. b, Across the metapopulation, patches in different successional stages are linked through seed rain. c, Evolutionary dynamics in the two traits give rise to a community with a characteristic pattern of coexisting species (white circles), comprising a few early-successional fast-growing species (green and blue) and a high diversity of late-successional shade-tolerant species (yellow to red). Colours in c correspond to those in a and b. In this schematic, patches are shown with finite size and spatial structure, whereas in the model they are considered to be large and spatially homogeneous.

The first trait we consider is the leaf-construction cost (LCC). LCC captures a tradeoff between the cost of building leaves and leaf turnover rate^15,20^ (Fig. 6a). Embedding this tradeoff in a physiological growth model gives rise to a demographic tradeoff between height growth rate and shade tolerance of seedlings (Fig. 6b). Correspondingly, LCC is regarded as one of several traits associated with successional status^21^. In our model, evolution in LCC leads to the emergence of three distinct successional types (Fig. 2a). Low LCC strategies correspond to early-successional types and flourish in high light conditions immediately following a disturbance. Due to their high leaf turnover, low LCC strategies are intrinsically shade-intolerant, and thus unable to regenerate in older patches, where higher LCC strategies prevail, corresponding to mid– and late-successional types. Our analysis is more realistic than earlier niche models as we explicitly link to physiological traits and account for size structure in light competition. Nevertheless, the results confirm core conclusions from earlier models: successional differences can only maintain a limited number of types differing in growth rate and shade tolerance, and only a single late-successional species^5–8^.

**Figure 2:**
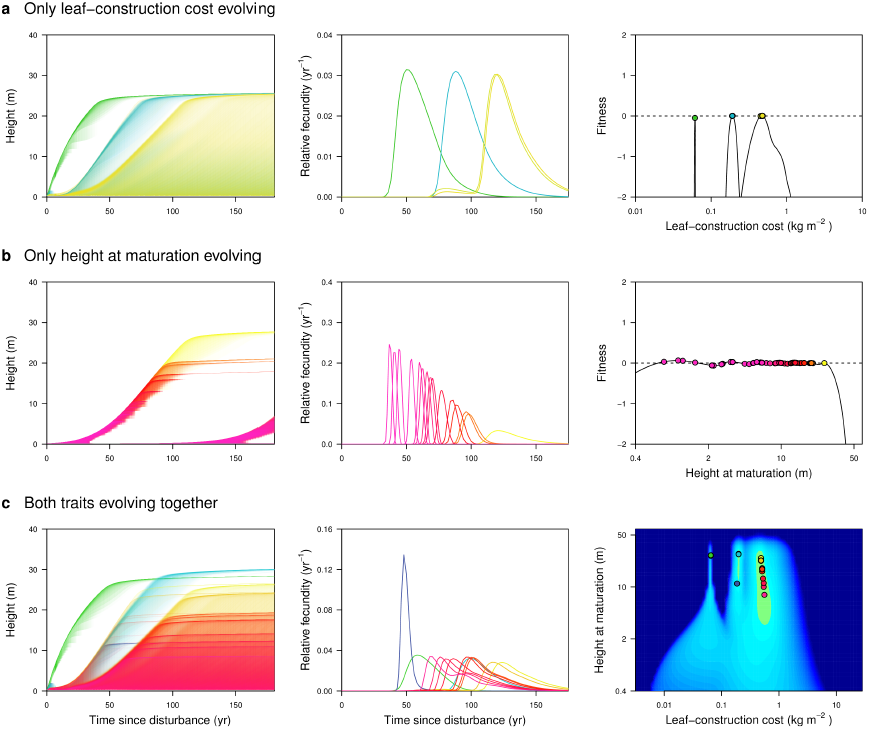
Multi-trait evolution under physiological tradeoffs offers alternative pathways of niche differentiation, leading to a concentration of late-successional types. Rows show communities resulting when **a** only leaf-construction cost evolves, **b** only height at maturation evolves, and **c** both traits evolve together. The first and second columns, respectively, show the size distributions and seed rains of species within a patch as it recovers from disturbance. Colours indicate trait values as in Fig. 1, while the intensity of shading in the first column indicates the density of individuals at a given height and patch age. The third column shows evolved trait mixtures and corresponding fitness landscapes. The fitness landscape in **c** is coloured according to Fig. 3. Only when the two traits evolve together, the resultant community structures match empirically observed patterns.

Surprisingly, very different conclusions emerge when a second trait interacts with LCC (Fig. 2b). Height at maturation (HMAT) is the height at which the allocation of surplus energy shifts from growth to reproduction (Fig. 7a). Changes in HMAT affect several demographic outcomes, including age at maturation, survival to maturation, and maximum fecundity (Fig. 7b). We find that differentiation in HMAT also promotes niche diversification, but through an entirely different mechanism than the one associated with LCC. Unlike LCC, differentiation in HMAT allows strategies with similar successional type (i.e., LCC) to coexist with one another via the partitioning of reproductive opportunity (Fig. 2b). As they have similar LCC, different HMAT strategies show the same patterns of seedling recruitment, growth rate, and shade tolerance. They differ in their growth rate at larger size due to differences in reproductive allocation. A series of strategies can coexist, with early maturing strategies being progressively over-topped by strategies with higher HMAT (Fig. 2b). Importantly, coexistence via this second mechanism is restricted to slow-growing high-LCC types, among which competitive thinning during stand development is less intense (Fig. 11).

Together, differentiation in these two traits can lead directly to a community showing the archetypal structure of most tropical forests^12,19^: a limited number of early-successional strategies, and a larger number of late-successional, shade-tolerant trees and shrubs (Figs. 2c). Such trait mixtures arise because the physiological tradeoffs considered here offer alternative pathways of niche differentiation, with interactive effects on diversity (Fig. 2). Previously, it was recognised that the widely cited competition-colonisation tradeoff used in many niche models may consist of two distinct tradeoffs, one involving differences in the ability of species to colonise vacant patches and another involving differences in the ability to grow at high versus low light^23^. Here, we have identified a third dimension of differentiation in the maturation schedule of larger plants with the same seedling demography. Moreover, we show that the particular conditions needed for diversification along multiple niche axes are widespread, if not universal, in forests: size-structured competition for light, a disturbance regime, and tradeoffs in plant function related to growth rate and reproductive allocation. Thus, the perception that niche models cannot explain *“why niches are more finely partitioned under low-light conditions than under high-light conditions”* ^12^ may have arisen only because previous niche models lacked critical features of real forests – most importantly, the ability for species to differentiate in traits influencing both seedling growth rate and reproductive allocation.

The two tradeoffs considered here do not only offer alternative pathways of niche differentiation. They also have fundamentally different effects on fitness landscapes, with implications for the total diversity of species that can be maintained. In general, a fitness landscape describes whether variants with new trait combinations can invade an existing resident population. Fig. 3a shows the fitness landscape of an empty community. Regions of positive fitness represent fundamental niche space: any trait combination within these regions can maintain a population, in the absence of competition with other types. As the community becomes more densely occupied, the fitness landscape is drawn downwards. Peaks, ridges, and valleys are created, and areas of trait space that still offer positive fitness (available niche space) become more restricted (Fig. 3b-h). Residents can draw down the fitness landscape at considerable distance from themselves in trait space, as well as nearby (Fig. 3). This reflects an important reality in plant communities: species with different traits nevertheless deplete the same basic resources. In our model, competition takes the form of reducing light availability, with impacts varying across patches of different age and individuals of different height. Interestingly, the eventual mixtures produced when evolving the two traits in combination (Fig. 3h) can be predicted neither from studying the evolution of each trait in isolation (Fig. 11), nor from scrutinizing the fundamental niche space (Fig. 3a).

**Figure 3:**
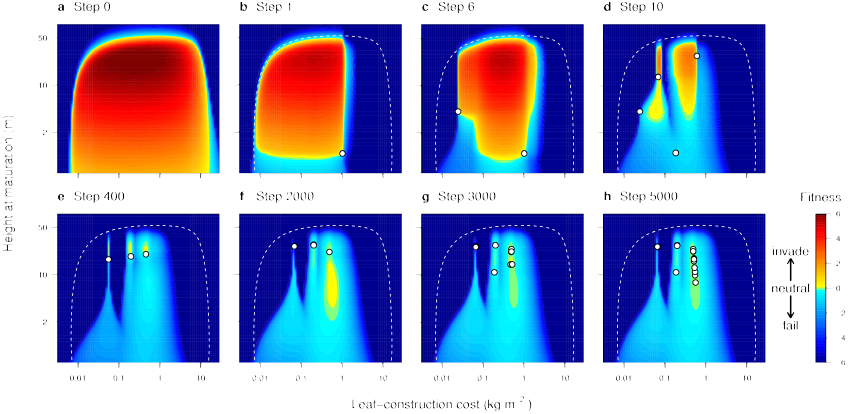
Community assembly leads to characteristically structured fitness landscapes, including peaks, valleys, and neutral ridges. Panels show fitness landscapes at different stages of assembly of the community shown in Fig. 1 & Fig. 2c. Colouring indicates the invasion fitness across trait space of rare species competing with the resident species (white circles). The fitness landscape guides the process of community assembly as only trait regions with positive fitness (coloured yellow to red, delineating the available niche space) can be invaded by new species. A ridge-shaped fitness plateau (green region in h) naturally emerges in the course of the assembly, enabling a high diversity of shade-tolerant plant species through evolved neutrality. For an animation spanning the entire assembly process, see Fig. 10.

Niche differentiation is traditionally thought of as creating a fitness landscape with distinct adaptive peaks, supporting a finite number of types that are able to persist without competitive exclusion^5,24^. Such a landscape indeed results when LCC is evolving alone (Fig. 2b; for pattern through assembly see Fig. 8). This expectation of spaced peaks in fitness landscapes underpins widespread use of limiting-similarity metrics in current plant ecology. Our model shows that fitness landscapes of quite different shape are also possible. When HMAT is evolving alone, it produces a flattened surface (Fig. 2b; for pattern through assembly see Fig. 9). When LCC and HMAT coevolve, a nearly level (near-neutral) plateau arises, in a specific region of trait space, corresponding to shade-tolerant late-successional types (Fig. 2c-3; for pattern through assembly see Fig. 10). Thus, deterministic niche differentiation can produce fitness landscapes not only with distinct peaks but also with flattened regions.

These plateaus in fitness landscapes give rise to what can be called ‘evolved neutrality’. They are far from causing the across-the-board neutrality (fitness equivalence) assumed by UNT^10,12^, which has widely been criticised as implausible in face of obvious functional differences^4,14^. Rather, they imply near-neutrality across tightly circumscribed zones of trait space that arise as outcomes of the competitive drawdown of fitness landscapes. In general, flat fitness landscapes slow competitive exclusion, and can thus support a much larger number of species than fitness landscapes that are coarsely rugged or possess spaced peaks. Hence, differentiation in height at maturation not only increases the number of types supported through entirely deterministic processes (compare Fig. 2a and Fig. 2b), but evolution in this trait ultimately also allows the accumulation of a large number of species with similar trait values via evolved neutrality.

So far we have focussed on the functional composition and diversity of tropical forests, because this is where previous niche models seemed least satisfying. Thus, the mixtures presented in Figs. 1-3 are assembled under conditions that can be thought of as akin to those in tropical rainforests: high productivity, and disturbance regimes with intermediate return intervals of around 50 yr^27^. We now expand our results from this focus: Fig. 4 shows how, by varying disturbance interval and site productivity, our model readily generates a much wider variety of trait mixtures and communities from the same basic set of processes. These evolved forests capture two well-known empirical patterns. First, LCC values are lower in sites of higher productivity and shorter disturbance interval^15,16,18^. Second, plants are generally taller in more productive sites with longer intervals between disturbances^16,17^. In addition, our model provides intuitive and testable hypotheses about other features of vegetation, for which good quantitative data are as yet lacking. First, the number of successional classes (LCC values) drops from three down to one as disturbances become more frequent, such that shade-tolerant types are absent in the sites with shortest intervals between disturbances (Fig. 12). Second, communities occurring on productive sites that are infrequently disturbed lack the diversity of late-successional species that arises at intermediate disturbance intervals, thus resembling tall temperate forests (Fig. 4e). Third, a pattern observed across all the evolved mixtures is that diversity is highest within the last successional type, irrespective of whether the community comprises one, two, or three successional types. As before, these diversity hotspots in trait space correspond to plateaus of evolved fitness equivalence (Fig. 12). Thus, while in tropical forests shade-tolerant late-successional types are diverse, and fast-growing early-successional types are not, in frequently disturbed communities (like fire-prone woodland or river banks) species diversity is predicted to shift to the fast-growing strategies (Fig. 4).

**Figure 4:**
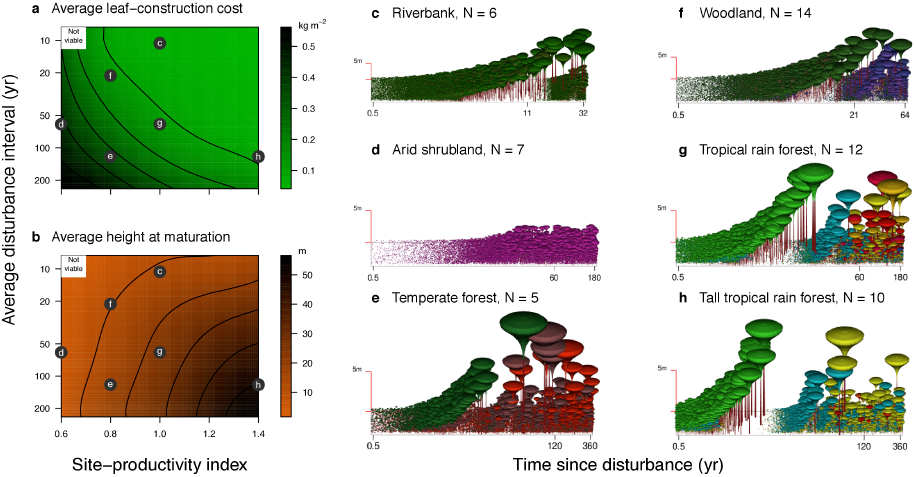
Extended niche models allow for natural variation of forest structure and diversity with environmental conditions, recovering broad empirical patterns. **a-b,** Mean trait values vary among communities evolved with different average disturbance intervals and site productivities. The resultant fitness landscapes are shown in Fig. 12. Letters refer to the communities illustrated in the subsequent panels. **c-h**, Resultant successional dynamics in a patch recovering from disturbance. The middle tick labels along the logarithmically scaled horizontal axes indicate the average disturbance intervals. Panel titles suggest real vegetation types bearing likeness to the model-evolved communities. Also shown are the numbers of distinct functional types in each community (determined as the number *N* of trait combinations with positive seed rain differing from one another by at least 1% in each trait after 5000 steps).

Overall, our work demonstrates that when a few key ingredients common to all forests are added, niche models can readily account for four empirically observed, ubiquitous, and thus fundamental, patterns of forest structure. The first is the long-term coexistence of species differing in growth rate and maximum size. Second is the tendency for trait values to shift along environmental gradients. Third is the feature of tropical forests that helped motivate the neutral theory: the relatively denser occupation of niche space in low light. Fourth is the remarkable species diversity of some forests, given that all plants compete for the same basic resources. To explain such coexistence, we have shown not only how a moderate diversity of species can coexist via niche differentiation, but also how niche differentiation produces trait regions of evolved neutrality wherein additional species will accumulate via neutral dynamics. Contributing to a growing body of work recognising the possibility of evolved (or emergent) neutrality as a way to reconcile niche and neutral theories of diversity^10,11,13,28^, our study provides the first mechanistic and calibrated example showing how evolved neutrality can arise. This provides a foundation for a synthetic theory of forest diversity, structure, and function.

## 3 METHODS SUMMARY

Our conclusions are based on a model in which site productivity, competition for light, and a disturbance regime drive the height distribution of individual plants within and among species (Figs. 1,5). A vegetation landscape is represented as a metapopulation of patches in different stages of development, linked by dispersal. Within patches, plants compete for light in accordance with their heights. Individuals of each species are thus organised along a continuous size axis, reflecting their heights, and the population is described by its height distribution.

**Figure 5:**
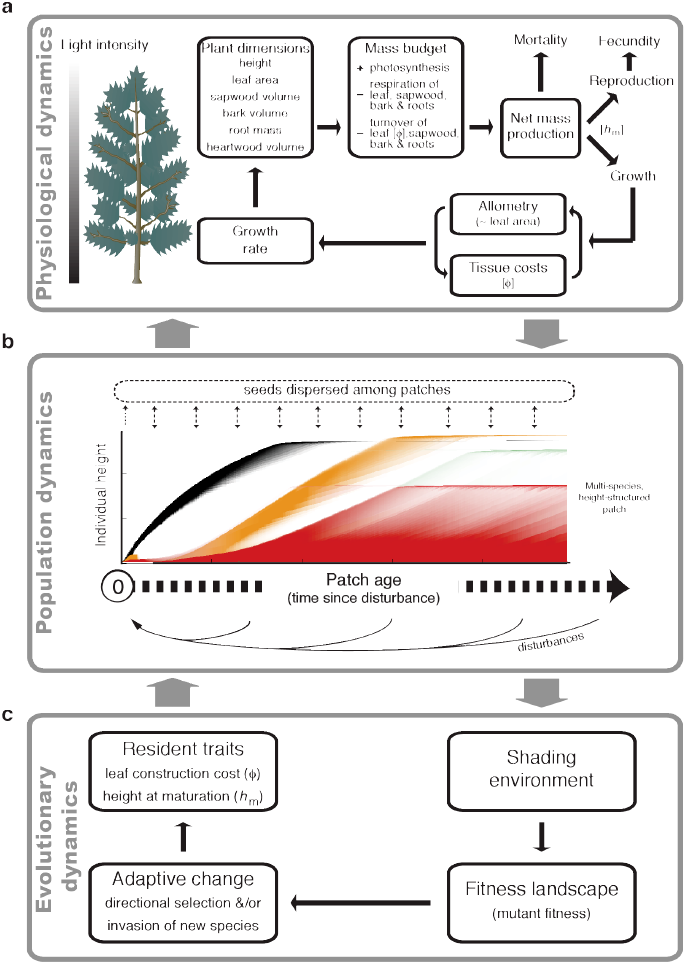
Overview of processes represented in the model, including physiological dynamics, population dynamics, and evolutionary dynamics. **a**An individual’s vital rates are jointly determined by its light environment, height, and traits. **b**, A metapopulation consists of a distribution of patches linked by seed dispersal. Disturbances occasionally remove all vegetation within a patch. Competitive hierarchies within developing patches are modelled by tracking the height distribution of individuals across multiple species (distinguished by colours) as patches age after a disturbance. The intensity of shading indicates the density of individuals at a given height. **c**, The traits of the resident species determine the shading environment across the metapopulation, which in turn determines fitness landscapes. Resident traits adjust through directional selection up the fitness landscape and through the introduction of new species where the fitness landscape is positive.

As patches age after a disturbance, their height distribution develops through the processes of growth, mortality, and reproduction. A detailed physiological model determines a plant’s growth, mortality, and fecundity rates from its height, traits, and light environment via net mass production (Methods).

The reproductive success of individuals in the model is frequency– and density-dependent. A new strategy can invade if, and only if, its net rate of population increase across the metapopulation (invasion fitness) is positive (Methods). Once established, resident strategies increase in abundance until they reach their carrying capacity; as a result, the invasion fitness of residents approaches zero at equilibrium.

Evolutionary analyses were focussed on deriving trait combinations favoured by the ecological model, and not on a realistic modelling of past evolution. New strategies were introduced into communities until no further significant evolutionary change occurred.

## 4 METHODS

### 4.1 Physiological dynamics

To describe the rates of growth, mortality, and fecundity, we used physiological functions developed in a previous publication^29^. These functions are specifically formulated for use in height-structured metapopulation models of the type employed here. Their principal ingredients are: (i) an allometric model linking an individual’s leaf area to its height and mass of supporting tissues, (ii) a continuous vertical distribution of leaf area within the crown of each plant, (iii) net dry-matter production given by gross photosynthesis (calculated by integrating instantaneous assimilation rates in a given light profile over the annual solar environment) minus costs of tissue respiration and turnover, (iv) fecundity calculated from reproductive investment and mass per seed, and (v) an exponential increase in mortality with declining carbon income per leaf area^29^.

The two evolving traits, leaf construction cost (LCC, *φ*) and height at maturation (HMAT, *h*_m_), influence growth, generating performance differences with respect to traits. The details of the trait-related tradeoffs are as follows. LCC determines the cost of deploying leaf area and is related to leaf turnover *k*_l_ according to a relationship observed in a global dataset of 678 species: *k*_l_ = 0.0286yr-1(*φ* m^2^kg^−1^ )^−171^ (Fig. 6). HMAT determines the fraction *r*(*h*_m_,*h*) of mass increment allocated to reproduction during development, varying with plant height *h:r*(*h*_m_, *h*) = 1/ (1 + exp(50(1 — *h/h*_m_))). This function describes a rapid transition from nearly 0% to almost 100% allocation to reproduction at *h = h*_m_ (Fig. 7). The remaining fraction 1 — *r*(*h*_m_, *h*) of mass increment is allocated to growth.

**Figure 6:**
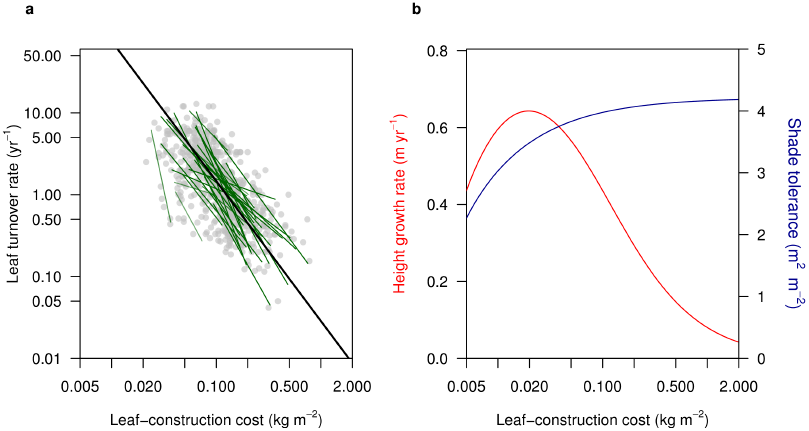
Trade-off between leaf-construction cost and leaf turnover rate, and its demographic consequences. a, Across species, leaf-construction cost (LCC) is inversely related to leaf turnover rate. Data are for 678 species from 51 sites15. Green lines show standardised major axes fitted to the data from each site, with the intensity of shading adjusted according to the strength of the relationship. The black line indicates the parameter values used in the current paper. **b**, Modelled height growth rate and shade tolerance for 25cm tall seedlings differing in leaf-construction cost. Shade tolerance is quantified as the amount of leaf area above the plant that can be endured before carbon budget becomes negative.

**Figure 7:**
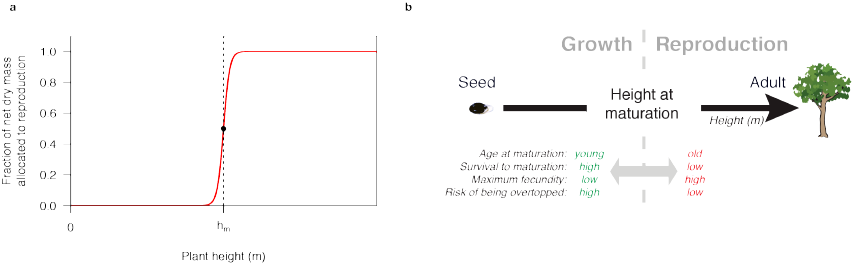
Trade-off between growth and reproduction, and its demographic consequences. a. The fraction of net dry mass produced allocated to reproduction varies throughout ontogeny, with a sharp increase from nearly 0% to nearly 100% when a plant reaches its height at maturation (HMAT). **b**, Height at maturation influences four key demographic factors, giving rise to several emergent life-history tradeoffs.

**Figure 8:**
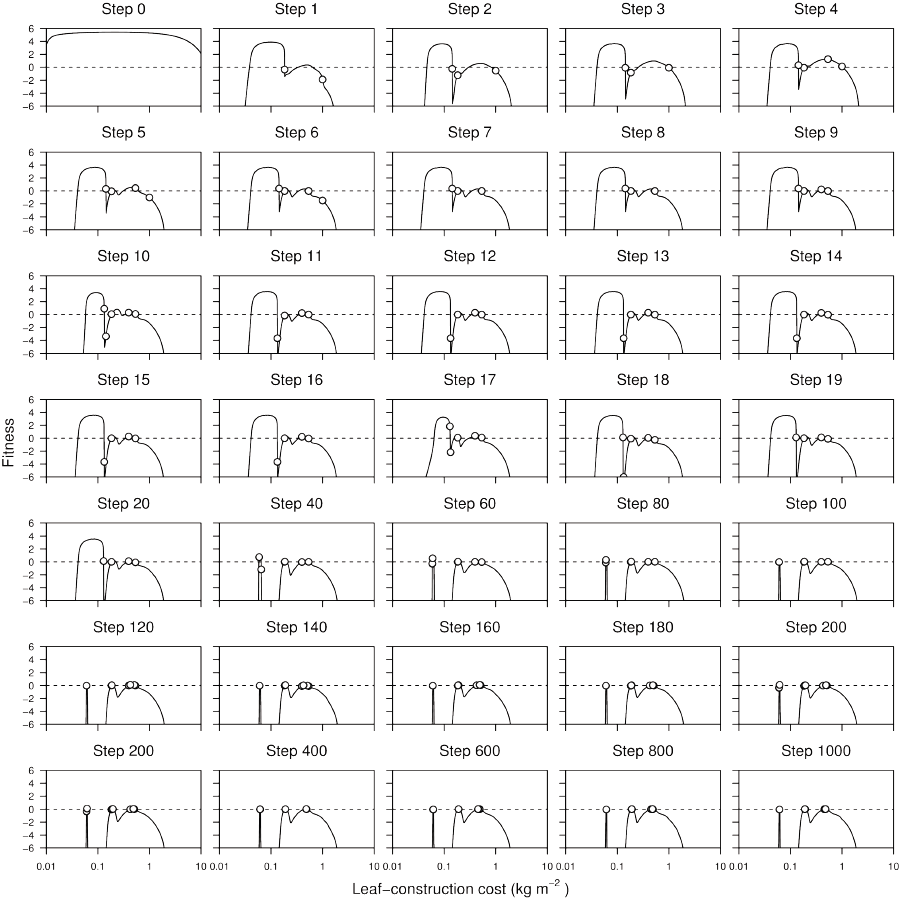
Community assembly when only leaf-construction cost is evolving. Panels show the resultant one-dimensional fitness landscapes at different moments of the assembly process for the community in Fig. 2a. Height at maturation was held constant at 23.5 m. See Fig. 12a for mixtures with other values of height at maturation.

**Figure 9:**
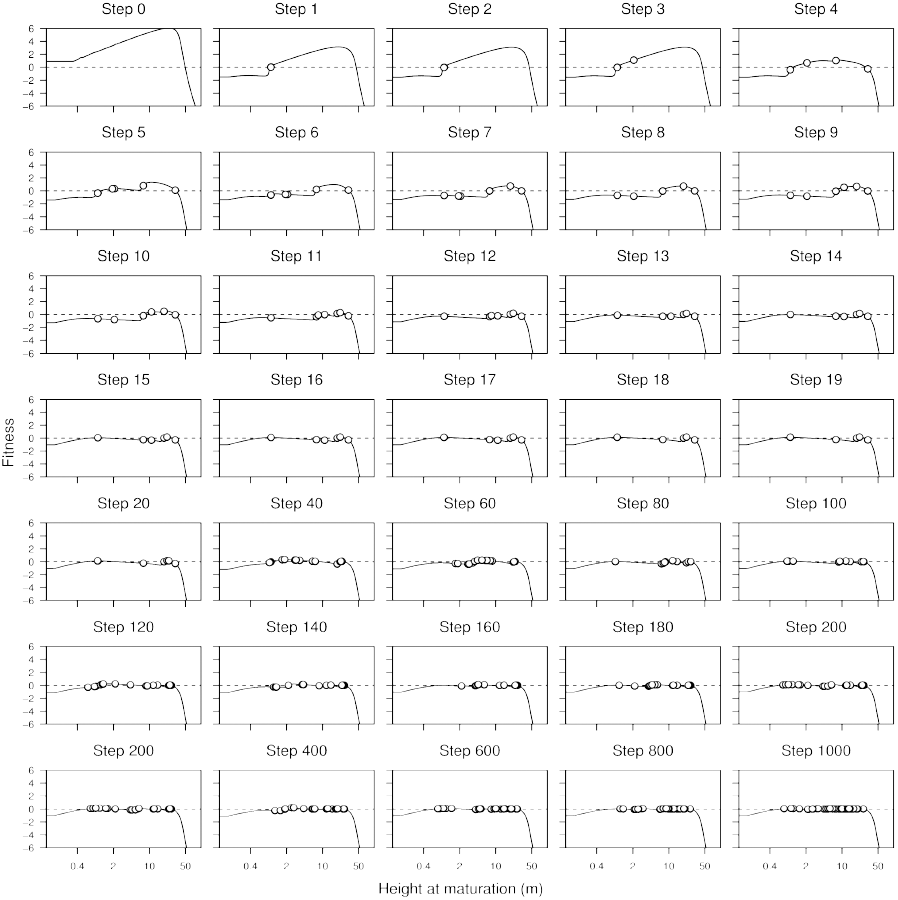
Community assembly when only height at maturation is evolving. Panels show the resultant one-dimensional fitness landscape at different moments of the assembly process for the community in Fig. 2b. Leaf-construction cost was held constant at 4.21 kg m-2. This value was chosen because it is situated within the region of parameter space where coexistence of different heights at maturation is possible (Fig. 11b). See Fig. 12b for mixtures with other values of leaf-construction cost.

**Figure 10:**
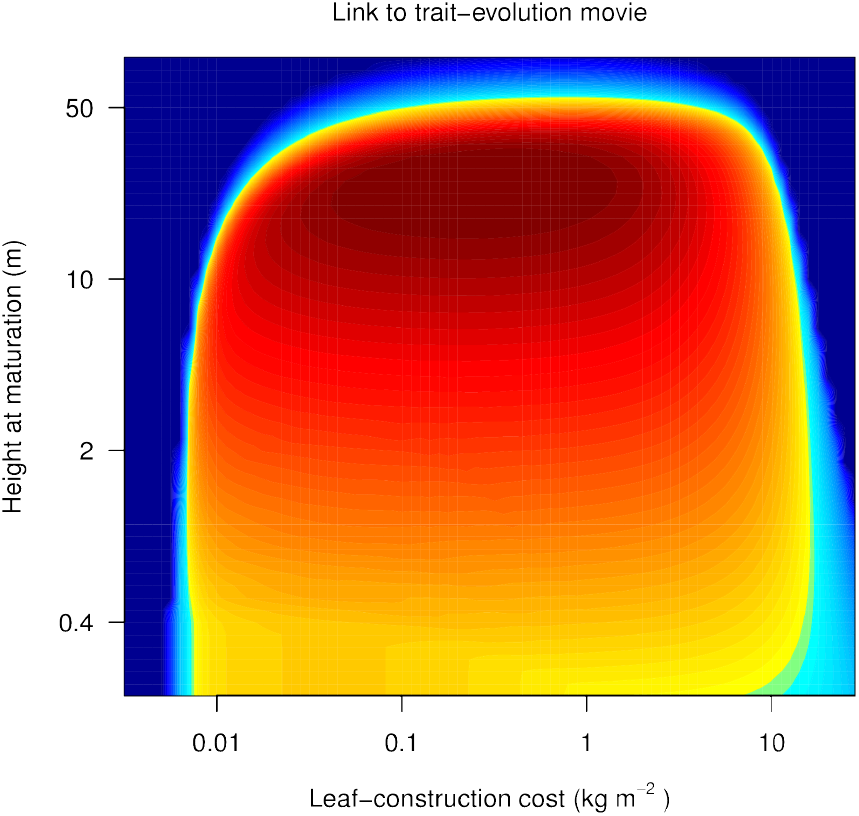
Animation of fitness landscape during bivariate trait evolution. This animation shows changes in the two-dimensional fitness landscape during the assembly process for the community shown in Figs. 1, 2c, and 3. Colouring indicates the invasion fitness across trait space of rare species competing with the resident species (white circles), as in Fig. 3

**Figure 11:**
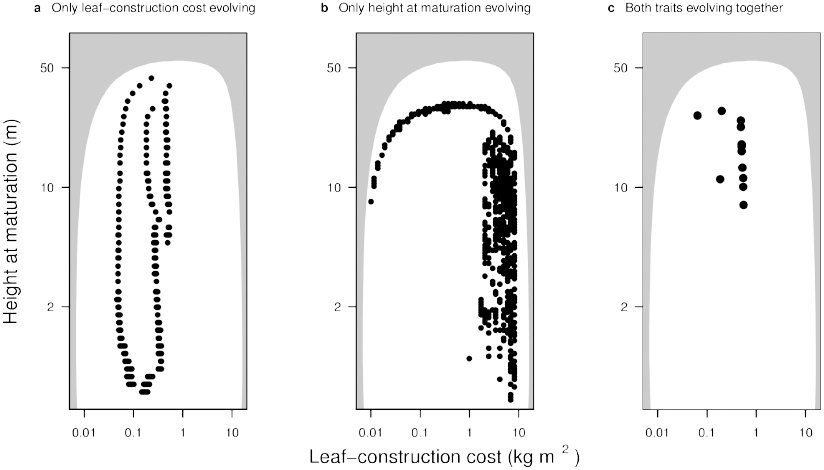
Rich interactions between the two traits determine bivariate trait mixture. Panels **a** and **b** shows the evolved univariate mixtures in each trait when the other trait is fixed at a specific value, while **c** shows the bivariate trait mixture that results when both traits are evolving together. The grey region in each panel indicates trait combinations that are not viable even in the absence of any competition (Fig. 3a). Note that in panel **b**, coexistence is only possible at high values of LCC.

**Figure 12:**
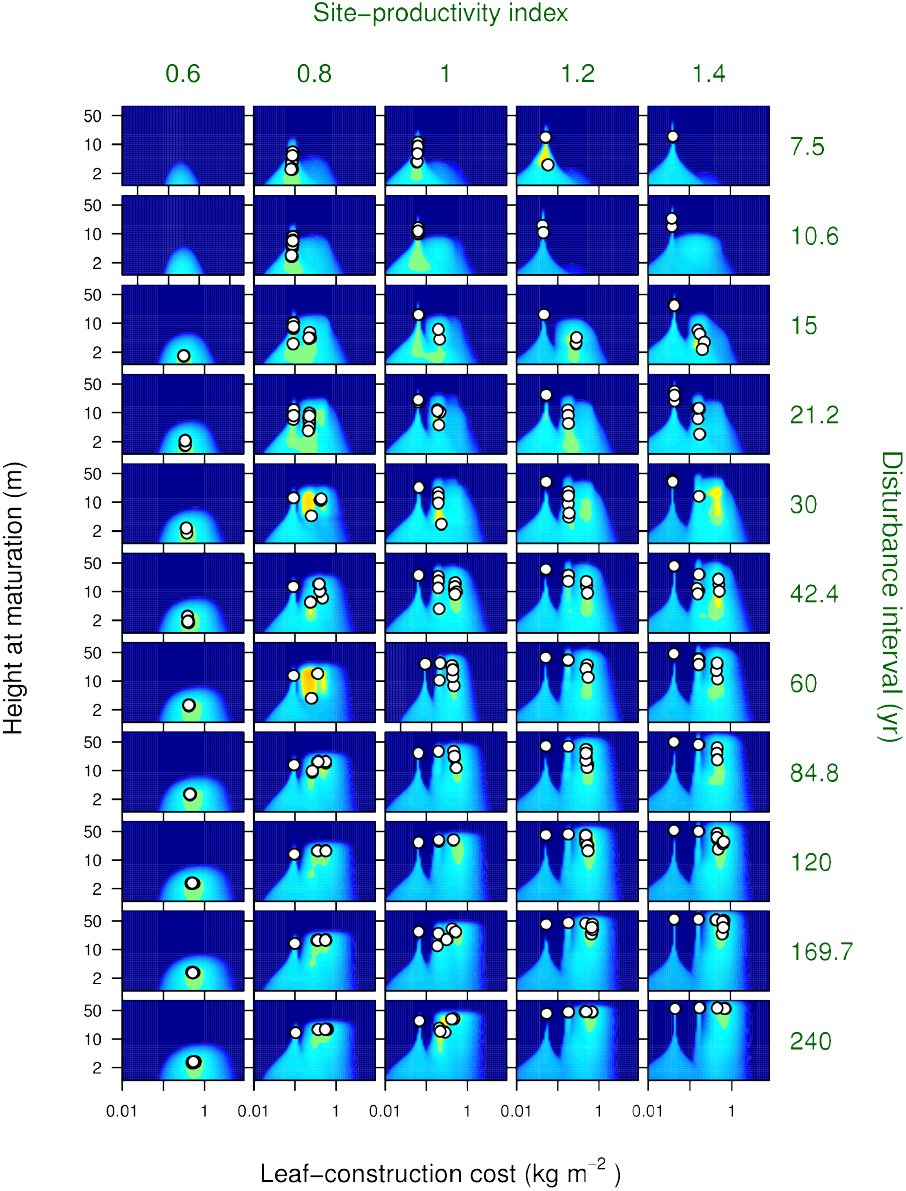
Effect of disturbance interval and site productivity on trait mixtures and fitness landscapes. Colouring indicates the invasion fitness across trait space of rare species competing with the resident species (white circles), as in Fig. 3 Plots show fitness landscapes after 5000 steps of the assembly process. After this time, most communities have reached a stable trait mixture. A few exhibit oscillatory dynamics in trait mixtures: above, these are distinguished by large residual regions of positive invasion fitness (yellow).

### 4.2 Population dynamics

We consider a metapopulation of patches, with the age *a* of each patch corresponding to the time since its last disturbance (Fig. 1). Disturbance events remove all individuals within a patch. Following a disturbance, patches are colonised by seeds arriving from a global pool of dispersers; any seeds produced within the patch also contribute to this pool. Different patches within the metapopulation are assumed to have similar abiotic properties, and thus differ only in their age. The state of a monomorphic metapopulation can therefore be characterized by the frequency distribution *p* of patch ages *a* and, for each patch age *a*, the height distribution of individuals within patches of that age.

The distribution *p* is determined by the disturbance regime. Here we assume that disturbances are caused externally, and can be described fully by the average time *â* between disturbances. Further, we let the age-dependent probability γ*(a)* of disturbance increase linearly with patch age *a*. Incorporating these assumptions into von Foerster’s equation for age-structured population dynamics^30^ leads to a Weibull distribution of disturbance intervals^31^ and the solution 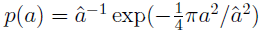.

Competitive hierarchies within developing patches are modelled by tracking the height distribution of plants as patches age after a disturbance. We denote by *x_i_* = (*φ_i_*, *h*_m._*_i_*) the traits of species *i*, where *φi* is the LCC and *h_m,i_* is the height at maturation, by *x* = (*x*_0_,… *x_N_*) the traits of *N* resident species, and by *y* = (*y*_0_,…,*y_N_*) their average seed rains across the metapopulation. Here traits refer to heritable characteristics of individuals that are constant throughout ontogeny, in contrast to an individual’s height, which changes through ontogeny. We denote by *n*(*x_i_*, *h*, *E_x,y,a_*) the density of individuals with traits *x_i_* and height *h* in a patch of age *a*, and by *g*(*x_i_*, *h*, *E_x,y,a_*), *d*(*x_i_*, *h*, *E_x,y,a_*), and *f*(*x_i_*, *h*, *E_x,y,a_*) the height growth rate, deathrate, and fecundity rate of those individuals. These rates depend on the vertical profile of shading within the patch, *E_x,y,a_*, which in turn depends on the traits *x* and the seed rains *y* of the resident types, as well as on the patch age *a*. For each trait *x_i_*, the seed rain *y_i_* is given by the cumulative seed output across the metapopulation,

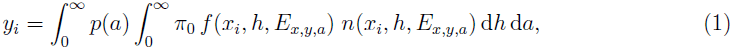

where π_0_ (a constant) is the probability of offspring surviving dispersal.

The dynamics of the plant community as described by the density distribution *n* are then given by the following partial differential equation^32–34^,

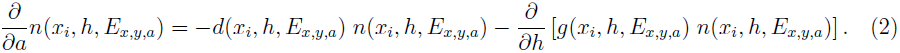

(Eq. 2 has two boundary conditions. The first,

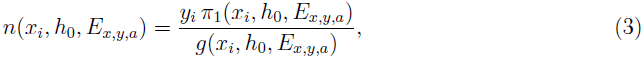

equates the inflow of individuals at the lower bound *h_0_* of the height distribution to the rate at which offspring are produced in the metapopulation. Here, *π_1_* is the probability of offspring successfully establishing themselves in a new patch, set to decrease with declining mass production, such that seedlings with zero or negative production fail to establish^29^. The second boundary condition states that patches are completely cleared after a disturbance,

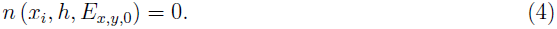

Individuals of all types interact via influences on a vertical shading profile, *E_x,y,a_*. Specifically, canopy openness at height *z* in a patch of age *a* is calculated from the cumulative leaf area above that height^35^,

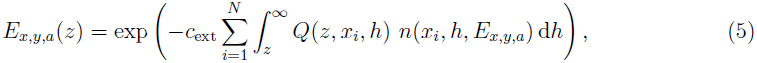

where *Q*(*z*, *x_i_*, *h*) is the total leaf area above height *z* for individuals of height *h* and traits *x_i_* and *c*_ext_ is the light-extinction coefficient.

### 4.3 Evolutionary dynamics

Communities were assembled based on stochastic mutations and immigration. At each step, we updated the population density of existing types according to their demographic fitness (see below). New types were introduced at a minimum density, and existing types that had dropped below the minimum density were removed. New types arose via two mechanisms: through small mutations in the trait values of existing residents, and through with traits values drawn from a uniform probability distribution across the entire viable trait space. The community-assembly process was continued for 5000 steps, after which the assembled communities were functionally fully differentiated. Although the community-assembly process can in principle be continuedbeyond the point of full functional differentiation, the results from our deterministic model are unlikely to be meaningful once random drift and environmental fluctuations would start to dominate the outcome in natural populations.

We denote by 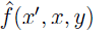 the (invasion) fitness of individuals with traits *x′* growing in a metapopulation with one or more resident strategies with traits *x* and seed rains *y*^36^. Invasion fitness is ideally calculated as the long-term per capita growth rate of the population with traits *x′* when rare; however, in structured metapopulations, the most convenient indicator of percapita growth rate is given by the logarithm of the basic reproduction ratio, 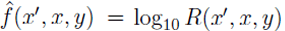^26^. Here, *R*(*x′*, *x*, *y*) describes the average number of dispersing seeds with traits *x′* produced by each dispersing seed with traits *x′* arriving in the metapopulation. Thus, the trait combination *x′* can invade when *R*(*x′*, *x*, *y*) > 1, which is equivalent to 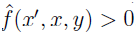.

Recalling that patches of age *a* have frequency density *p*(*a*) in the landscape, it follows that a dispersing seed with traits *x′* has probability *p(a)da* of landing in a patch of age *a* to *a* + da. The basic reproduction ratio for individuals with traits *x′* is thus given by

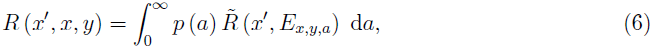

where 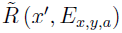 is the expected number of dispersing seeds with traits *x′* produced by a single dispersing seed with traits *x′* arriving in a patch of age *a*^26^. 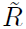 is calculated by integrating an individual’s fecundity over the expected lifetime of the patch, taking into account competitive shading from residents with traits x, the individual’s probability of surviving, and its traits,

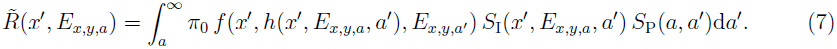

Here,

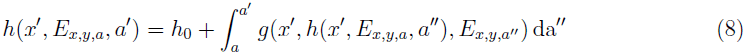

and

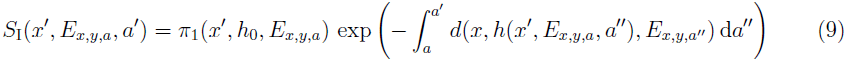

are the height and survival probability of dispersing seeds with traits *x* that arrive in a patch of age *a* at age *a*, while

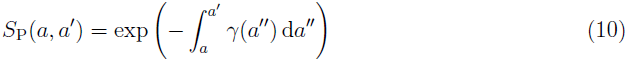

is the probability that the patch remains undisturbed from age *a* to *a′*.

### 4.4 Numerical analysis

To solve Eq. 2, we use a refined version of the Escalator Boxcar Train method^32^ powered by an embedded fourth-order Runge-Kutta ordinary differential equation (ODE) solver^37^. The ODE solver is also used to calculate invasion fitness by numerically integrating Eq. 7-10.

Unless otherwise specified, we use the parameter values described in the previously published growth model^29^. Our results are robust to changes in these parameters: Figs. 13 anf 14 show how such changes only slightly affect, quantitative details of the evolved bivariate trait mixtures, while leaving the qualitative features of the reported coexistence patterns unaffected.

**Figure 13:**
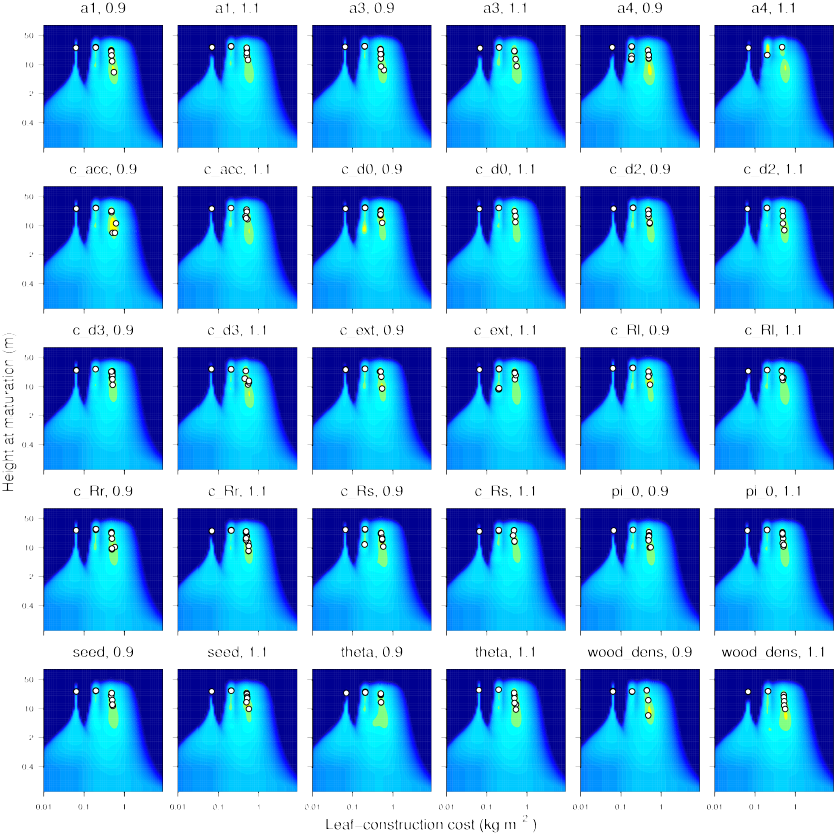
Sensitivity to changes in a range of model parameters. Each plot shows the community obtained when a single parameter is changed to 0.9 or 1.1 times its original value. Colouring indicates the invasion fitness across trait space of rare species competing with the resident species (white circles), as in Fig. 3 Plots show fitness landscapes after 5000 steps of the assembly process.

**Figure 14:**
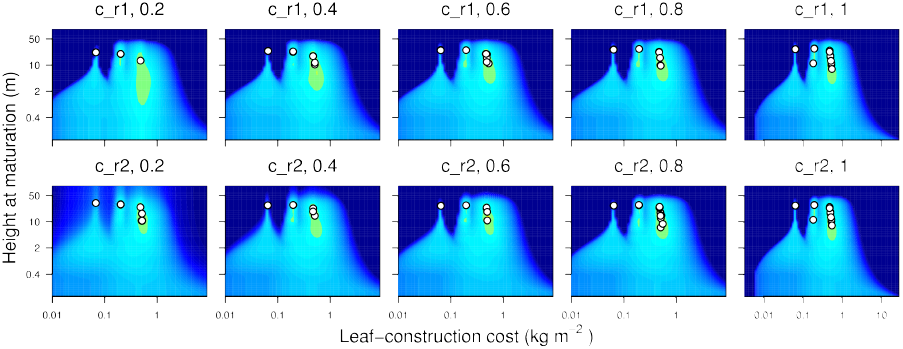
Sensitivity to changes in the shape of the reproductive allocation function. Each plot shows the community obtained when a single parameter determining the shape of the reproductive allocation function is changed to a fraction of its original value. Colouring indicates the invasion fitness across trait space of rare species competing with the resident species (white circles), as in Fig. 3 Plots show fitness landscapes after 5000 steps of the assembly process.

